# The nuclear receptor DHR3/Hr46 is required in the blood brain barrier of mature males for courtship

**DOI:** 10.1101/2021.03.30.437714

**Authors:** Chamala Lama, Cameron R. Love, Hoa Nhu Le, Jyoti Lama, Brigitte Dauwalder

**Affiliations:** Department of Biology and Biochemistry, University of Houston, Houston, TX, United States

## Abstract

The blood brain barrier (BBB) forms a stringent barrier that protects the brain from components in the circulation that could interfere with neuronal function. At the same time, the BBB enables selective transport of critical nutrients and other chemicals to the brain. Many of these processes are still poorly understood. Beyond these functions, another recently recognized function is even less characterized, specifically the role of the BBB in modulating behavior by affecting neuronal function in a sex dependent manner. Notably, signaling in the adult *Drosophila* BBB is required for normal male courtship behavior. Courtship regulation also relies on male-specific molecules in the BBB. Our previous studies have demonstrated that adult feminization of these cells in males significantly lowered courtship. Here, we conducted microarray analysis of BBB cells isolated from males and females. Findings revealed that these cells contain male- and female-enriched transcripts, respectively. Among these transcripts, nuclear receptor DHR3/Hr46 was identified as a male-enriched BBB transcript. DHR3/Hr46 is best known for its essential roles in the ecdysone response during development and metamorphosis. In this study, we demonstrate that DR3/Hr46 is specifically required in the BBB cells of mature males for courtship behavior. The protein is localized in the nuclei of sub-perineurial glial cells (SPG), indicating that it might act as a transcriptional regulator. These data provide a catalogue of sexually dimorphic BBB transcripts and demonstrate a physiological adult role for the nuclear receptor DH3/Hr46 in the regulation of male courtship, a novel function that is independent of its developmental role.

**Author summary:** The blood brain barrier very tightly regulates which molecules can enter the brain. This is an important protection for the brain, however, it also complicates communication between molecules in the circulating fluid and the brain. In fly courtship, for example, circulating male-specific products are crucially required for normal courtship. But the neuronal circuits that ultimately control the behavior are inside the brain, separated from these molecules by the blood brain barrier. The mechanisms of this communication are not known. Here we show that the blood brain barrier itself contains sex-specific RNAs and we show that one of them, a nuclear receptor called DHR3, is required in adult males for normal courtship. These findings promise new insight into the communication between blood brain barrier and the brain.

## Introduction

It is well established that the two layers of glial cells that tightly surround the nervous system form the *Drosophila* blood brain barrier (BBB)(1). Flies have a non-vascular open circulatory system that distributes the hemolymph. The BBB forms the tight exclusion barrier that is essential to protecting neurons from hemolymph components that could interfere with neuronal function (2, 3). At the same time, the barrier needs to allow selective uptake of nutrients and other molecules needed for brain function. The *Drosophila* blood brain barrier (BBB) surrounds the brain like a tight cap. It consists of two layers of glial cells. The outer perineurial glia cells (PG cells) are thought to function as a barrier for large-molecular weight molecules. The inner layer, the subperineurial glia (SPG), is adjacent to the neuronal cell bodies and contains the tight junctions that form the physical barrier (Fig 1A). It has been shown in a number of genetic and functional studies that the barriers in flies and vertebrates share not only structure and function, but also many homologous proteins that ensure their function, as shown in (4). A recent microarray study of isolated BBB cells has expanded on these earlier findings and shown that besides the characteristic barrier proteins, fly and mouse BBB cells share a large number of conserved proteins (5). That study has also provided the first detailed “inventory” of these cells in *Drosophila.* While the barrier and selective uptake functions of the BBB are its most obvious essential function, evidence is starting to accumulate that other physiological processes in BBB cells are contributing to brain function. For example, the G-protein-coupled receptor *moody* is specifically expressed in the subperineurial glial cells (SPG)(6, 7). While the absence of both *moody* isoforms leads to a leaky BBB (6, 7), mutants with only one of the isoforms have intact barriers, but behavioral defects in their response to cocaine and alcohol (6). In addition, *moody*, in a function independent of its function in barrier integrity, is also required in BBB cells for normal male courtship (8). That active signaling processes in BBB cells regulate neuronal output was further indicated by the finding that BBB-specific reductions in the G protein Galpha(o) cause courtship defects, while leaving the barrier integrity intact (8). It has been found that the circulating hemolymph contains male-specific factors from the fat body that are needed to ensure normal courtship (9). It is not clear how these factors interact with the male brain circuits that regulate the behavior. Here we examine whether the BBB expresses sexspecific transcripts that might be part of this communication. This would be in agreement with the finding that feminization of the BBB by expression of the feminizing TRA protein specifically in the BBB of adult males results in a significant reduction in male courtship (8). In these experiments, the tightness of the barrier was unaffected, suggesting that specific male transcripts are physiologically participating in courtship control. The identity of these factors and their function is unknown. Here we identify sex-preferentially expressed transcripts in the BBB of males and females and demonstrate that the nuclear receptor DHR3/Hr46, best known for its roles in larval development (10–12), is physiologically required in the BBB of adult mature males to ensure normal male courtship behavior.

**Figure 1.**
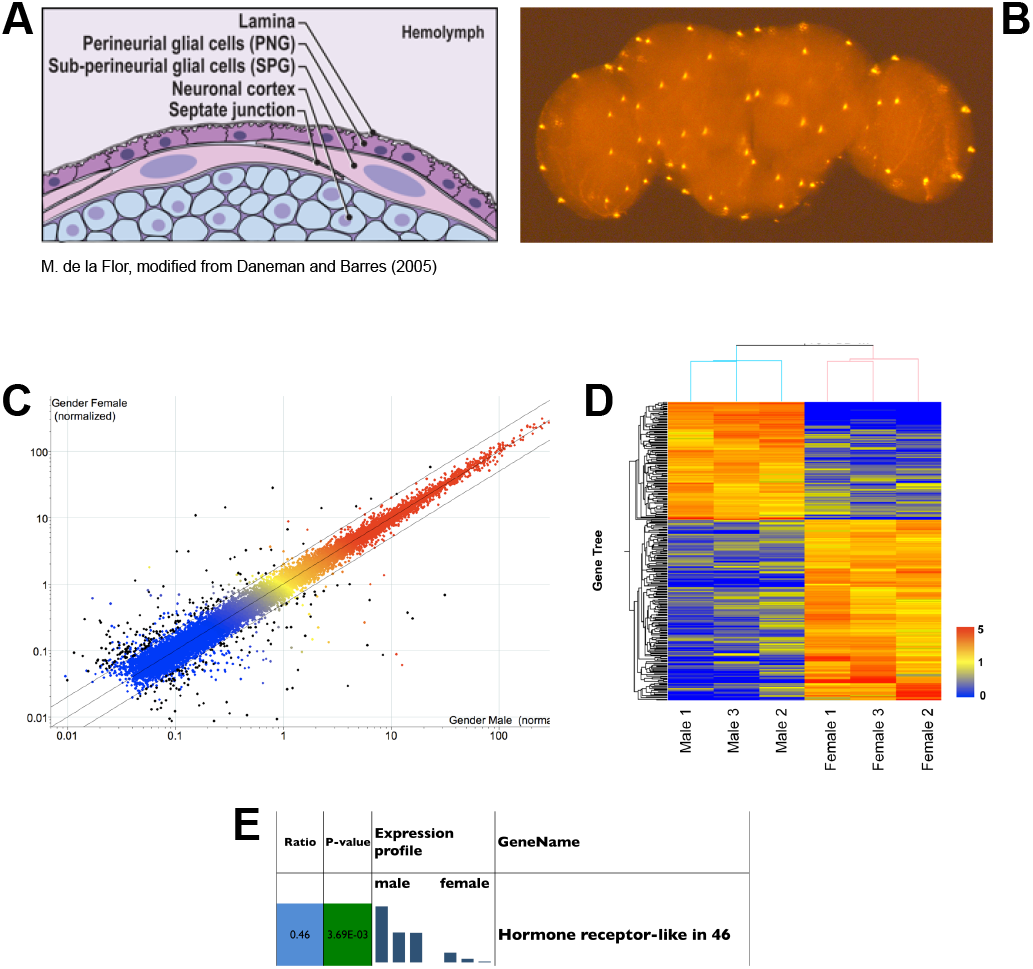
Microarray analysis of isolated SPG cells of the BBB. (A) Schematic of the *Drosophila* Blood Brain Barrier (BBB) (modified from (37). The BBB consists of two layers of glial cells, the outer Perineurial Glia (PG) facing the circulating hemolymph, and the inner Subperineurial Glia (SPG) with septate junctions that form the main barrier. The SPG is in contact with the underlying nuclei of the neuronal cortex. (B) Isolated fly brain with SPG cells labeled by nuclear DsRed expression driven by *SPG-Gal4.* Fluorescently marked cells like these from males and females were hand-isolated for RNA extraction. (C) Probes present (above background) in all male or female samples are displayed as normalized to the 75th percentile intensity of each array (19,218 probes). Each spot is the mean of 3 samples from each condition. Black spots =differentially expressed genes (>2Fold, T-test p-value < 0.05, 284 probes). Red/orange=High expression, Yellow=Medium expression, Blue=Low expression. (D) Differentially expressed genes (>2 fold,T-test p-value < 0.05) in Male vs. Female are displayed as normalized to the median value of each probe across six samples (284 probes). The heat map color scale is shown on the right. (E) DHR3/Hr46) is preferentially expressed in males.

## Results

### A microarray screen identifies male- and female-enriched transcripts in the BBB

In our previous experiments, there was a strong reduction in male courtship when we conditionally feminized adult BBB cells (8). This suggests that feminization disrupts male-specific transcripts that are physiologically required for normal mating behavior. In order to identify these transcripts, we isolated BBB cells from males and females and characterized their transcripts. The Gal4/UAS system was used to mark these cells (13). We expressed the fluorescent protein DsRed in the nuclei of SPG cells, using the *moody-Gal4* driver that drives expression in SPG cells *(SPG-Gal4)* (6). As seen in Fig 1B, the large nuclei of the SPG cells were specifically marked. We dissected fly brains and manually removed and collected fluorescent cells. Cells from approximately 50 flies were pooled for each biological replicate, and the RNA of three biological replicas from separate crosses was prepared for each sex. The RNA was subsequently used for microarray analysis by GenUs Biosystems (http://www.genusbiosystems.com/). The results confirmed the presence of sex-preferentially expressed transcripts in the BBB of males and females, respectively. 284 transcripts were identified that were enriched > 2 fold in either males and females (Figures 2C, D). Of those, 112 were male-preferentially expressed (S1 Table). As expected, the male-specifically expressed *rox* RNAs that are required for dosage compensation were highly specific to males. Furthermore, sexspecific *dsx* transcripts were identified because male and female transcripts use different polyA-sites and can thus be identified by microarray (14). An analysis of the GeneOntology of the enriched transcripts is shown in Table S2. Sex specific categories such as dosage compensation and sex determination are well represented, further confirming that the isolated cells are sexually determined. About half of the genes fall into one of these categories. The rest of the genes could not be assigned to a specific category. In addition to identifying sex-preferentially expressed RNAS, the experiment also provided an inventory of RNAs present in the BBB cells. The vast majority of BBB transcripts is equally expressed in males and females. Among them, as predicted for SPG cells, were RNAs that are characteristic of BBB cells (4, 5): RNAs for the junction proteins *sinu* and *neurexin,* for example, and the previously characterized SPG transcripts for *moody* and *Mdr65.* The most likely contaminating cells from the dissections would be fat body cells which are in close proximity to the BBB, and neuronal cells. We found very small amounts of the fat body transcript *Lsp-2,* or of the neuronal marker *elav.* They were not preferentially present in males or females, indicating that low amounts of these cells are unlikely to affect the identification of sex-specific transcripts in the BBB.

**Figure 2.**
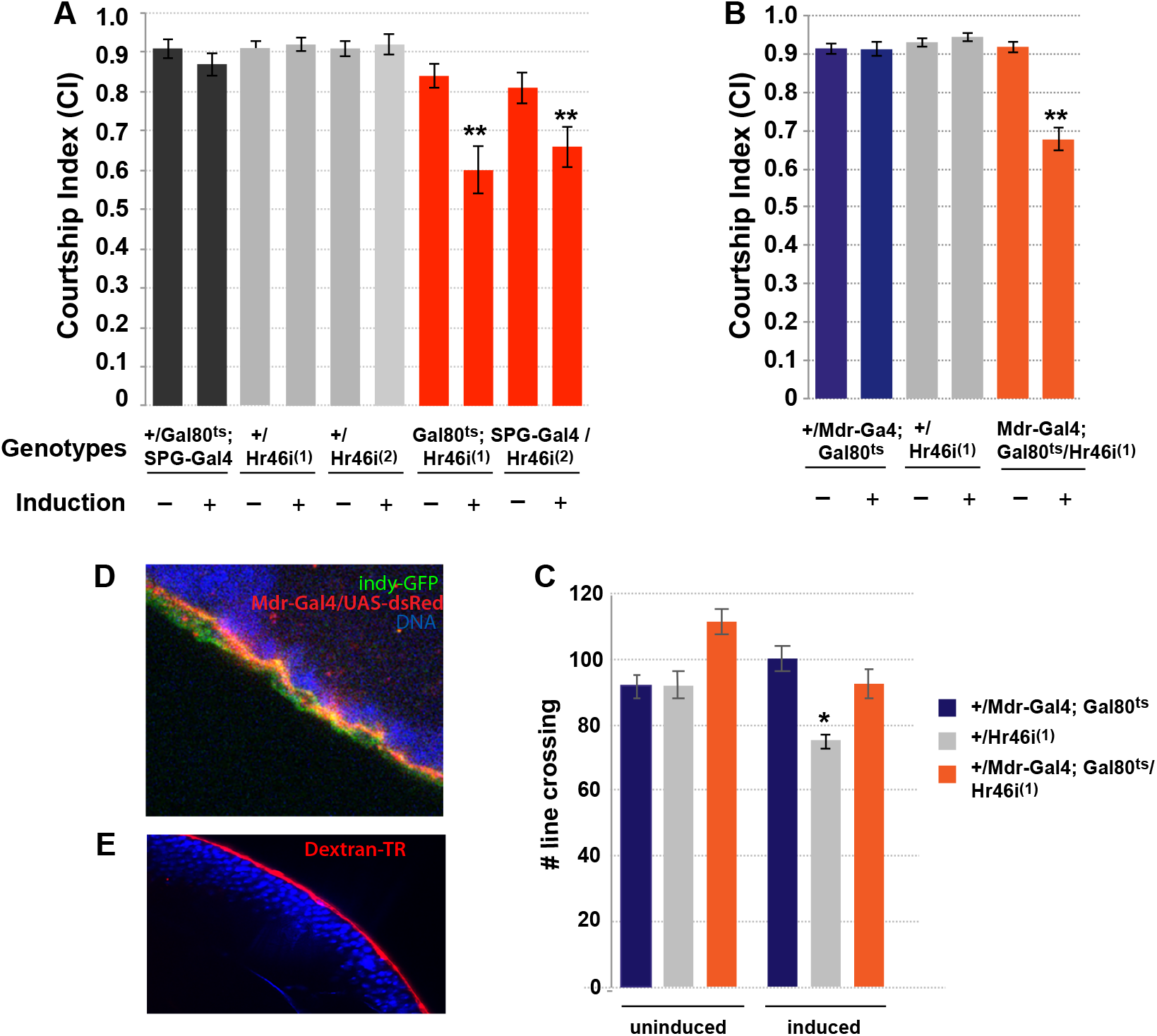
Knockdown of DHR3/Hr46 in the BBB of mature males reduces courtship. Graphs show the courtship index CI (fraction of time males spend courting during the observation period) ± SEM (A, B), or the performance of males in a control activity assay (# of line crossings ± SEM) (C) of the indicated genotypes. N= 20. Data were analyzed by ANOVA followed by Tukey multiple comparisons (p<0.05). Indices that are significantly different from the controls are marked by asterisks. *UAS-Hr46i* expression is restricted by the presence of *tubP-Gal80^ts^* at 18°C (induction -). Placement of 5-day-old males at 32°C for 16-24 hours (induction +) releases the Gal80 inhibition and leads to the expression of RNAi. (A) Expression of two different *UAS-Hr46-RNAi* (1-line 27253 and 2-line 27254) using *SPG-Gal4* significantly reduces male courtship. B) Conditional expression of *UAS-Hr46i* (27254) using *Mdr-Gal4* in adult males similarly reduces courtship in comparison to controls. The controls are 1) *+/Gal80^ts^;+/SPG-Gal4* and *+/Mdr-Gal4; +/ Gal80^ts^*, respectively and *2*) *+; +/UAS-Hr46 RNAi.* (C) The activity of the mutants as measured by number of line crossings is not lower than in control flies. *+/Hr46*^(1)^ control flies have reduced activity after induction that does not correlate with their courtship index. (D) *Mdr-Gal4* expression in SPG cells visualized by expression of *UAS-dsRed* in dissected brain (red). For comparison, *indy-GFP* (green) is expressed in both PG and SPG cells. (E) Blood-brain barrier integrity is not compromised in *Mdr-Gal4/ UAS-Hr46-RNAi* males. Flies were injected with 10 kDa TR-Dextran (red) and dye penetration into or exclusion from the brain was examined by confocal microscopy. The brain nuclei are stained with DAPI. A tight BBB is indicated by the demarcated red line on the surface of the brain indicating exclusion of TR-dextran from the brain of *Mdr-Gal4/ UAS-Hr46-RNAi* males.

### The nuclear receptor DH3/Hr46 is required in the BBB for courtship

One of the male-enriched BBB RNAs is the transcript for the nuclear receptor *DHR3/Hr46* (Fig 1E). DHR3/Hr46 is an orphan nuclear receptor that is most related to the mammalian ROR receptor (Retinoic acid related orphan receptor). DHR3/Hr46 is a well-described transcriptional regulator of larval developmental processes in response to ecdysone, but no adult functions have been described so far. To examine whether *DHR3/Hr46* is required in the BBB for courtship we conditionally expressed two different *DHR3/Hr46-RNAi* constructs specifically in the BBB of mature adult males and examined their courtship (Fig 2). Male courtship in *Drosophila melanogaster* consists of well-defined stereotyped behavioral steps that can easily be quantified in a courtship index (CI) (15–17). The CI is calculated as the fraction of time the male spends displaying any element of courtship behavior (orienting, following, wing extension, licking, attempted copulation, copulation) within a 10 minute observation period (18). We used the *Gal4/Gal80^ts^*system to restrict knockdown to mature males (19). Two different *BBB-Gal4* drivers were used to direct expression, the previously described *SPG-Gal4,* and a SPG-cell-specific *Mdr65-Gal4* driver that was generated in our lab (Fig 2D). The ATP binding cassette (ABC) transporter *Mdr65* has been shown to be specifically expressed in the SPG cells of the BBB (20, 21). Control flies containing a copy of just one of the two respective transgenes were grown, treated and tested in parallel to the knockdown flies as controls. At 18°C, Gal4 is inhibited by Gal80^ts^, and *DHR3/Hr46-RNAi* is not expressed. At this temperature, all genotypes exhibited normal courtship. In contrast, following induction at 32°C, males expressing *DHR3/Hr46-RNAi* in the BBB had significantly reduced courtship (p≤0.001) (Figs 2A, B). Reduction was observed with both drivers in combination with either of two different *UAS-DHR3/Hr46-RNAi* constructs. While courtship was reduced, the males were capable of performing all of the steps of courtship, but they did so with lower probability. To eliminate general sickness of the males as a cause for the reduced courtship, we performed a short-term activity assay (22) and found no activity defects in the knockdown flies (Fig 2C). We conclude that *DHR3/Hr46* is required in the BBB of mature males for normal courtship behavior. SPG BBB cells are glial cells. To confirm the glial requirement for DHR3/Hr46 we used the glia-specific driver *repo-Gal4* to drive *UAS-DHR3/Hr46-RNAi* in adult males and observed equally reduced courtship (p<0.001) (Fig 3A). As expected, when we expressed *DHR3/Hr46-RNAi* in the BBB with *Mdr-Gal4* in the presence of a glial-expressed Gal80 blocker *(repo-Gal80)* (23) we observed a reversal of the courtship defects. Together our findings demonstrate that *DHR3/Hr46* is needed in the glial SPG cells for normal courtship.

**Figure 3.**
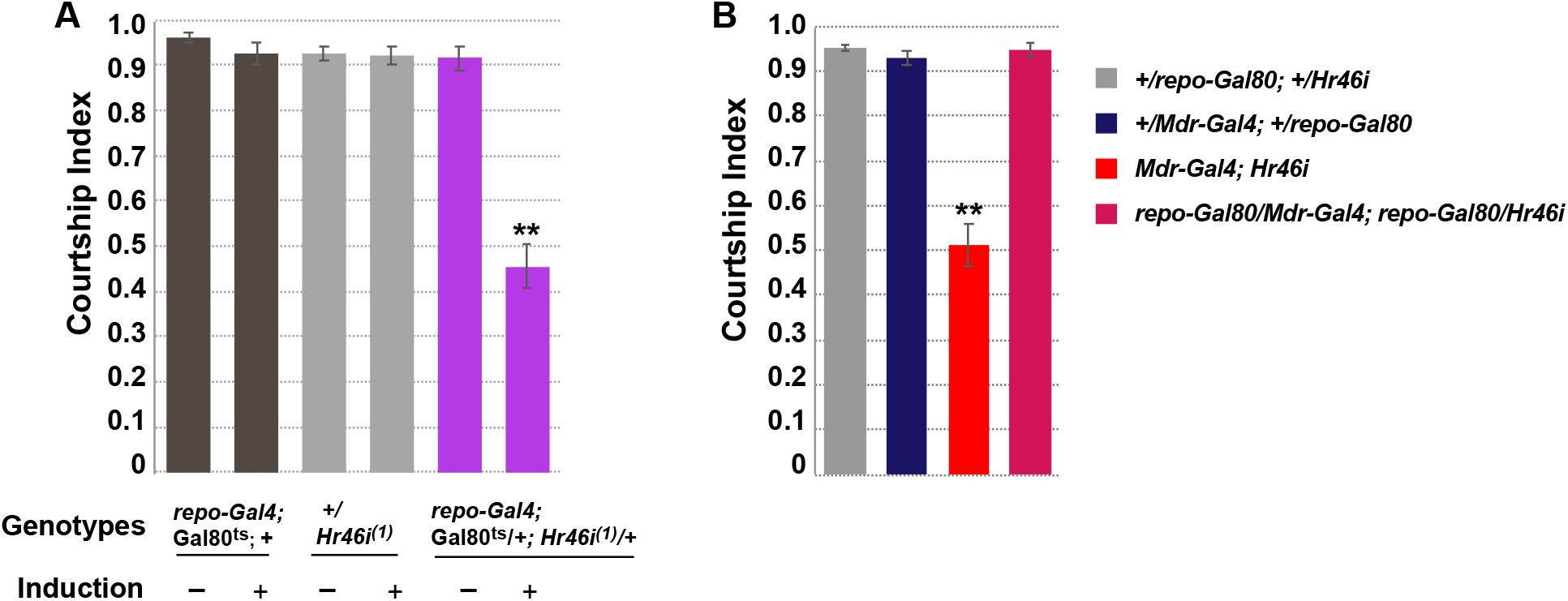
Hr46/DHR3 is required in the glial SPG cells for courtship. Graphs show the courtship index CI (fraction of time males spend courting during the observation period) ± SEM. N= 20. Data were analyzed by ANOVA followed by Tukey multiple comparisons (p<0.05). Indices that are significantly different from the controls are marked by asterisks. (A) Conditional glial knockdown of *DHR3/Hr46* in mature males using *repo-Gal4; Gal80^ts^* reduces courtship. (B) The courtship reduction of *Mdr-Gal4* directed *DHR3/Hr46* knockdown can be reversed by Gal80 expression in glial cells directed by *repo-Gal80.*

To assess whether *DHR3/Hr46* knockdown affects the integrity of the BBB, we tested the tightness of the BBB by injecting 10kD Texas-Red (TR)-marked Dextran. It is well documented that in wildtype animals TR-Dextran will be kept out of the brain and accumulate at the BBB, whereas a leaky BBB would allow entry into the brain (6). As shown in Fig 3E, males expressing *DHR3/Hr46* RNAi have normal BBB barrier function with the dye accumulating at the barrier. These findings indicate that BBB integrity is not compromised in the mutants, giving support to a physiological function for *DHR3/Hr46* in the BBB that is required for normal regulation of courtship.

### DHR3/Hr46 and its ligand are present in SPG nuclei

To examine the intracellular distribution of DHR3/Hr46, we used antibodies generated by the Thummel lab (11) to study the protein distribution in SPG cells of mature animals (Fig 4). Indy-GFP was used as a BBB marker; it is expressed in both PG and SPG cells (5). DNA was labeled with DAPI. BBB cells are big flat cells with large polyploid nuclei (24). Anti-DHR3/Hr46 antibody staining detected DHR3/Hr46Hr46 in the cytoplasm and the nucleus of SPG cells (Figs 4 A-D). DHR3/Hr46 is a transcriptional activator in larvae, and its presence in the nucleus of BBB cells is consistent with a transcriptional role in these cells.

**Figure 4.**
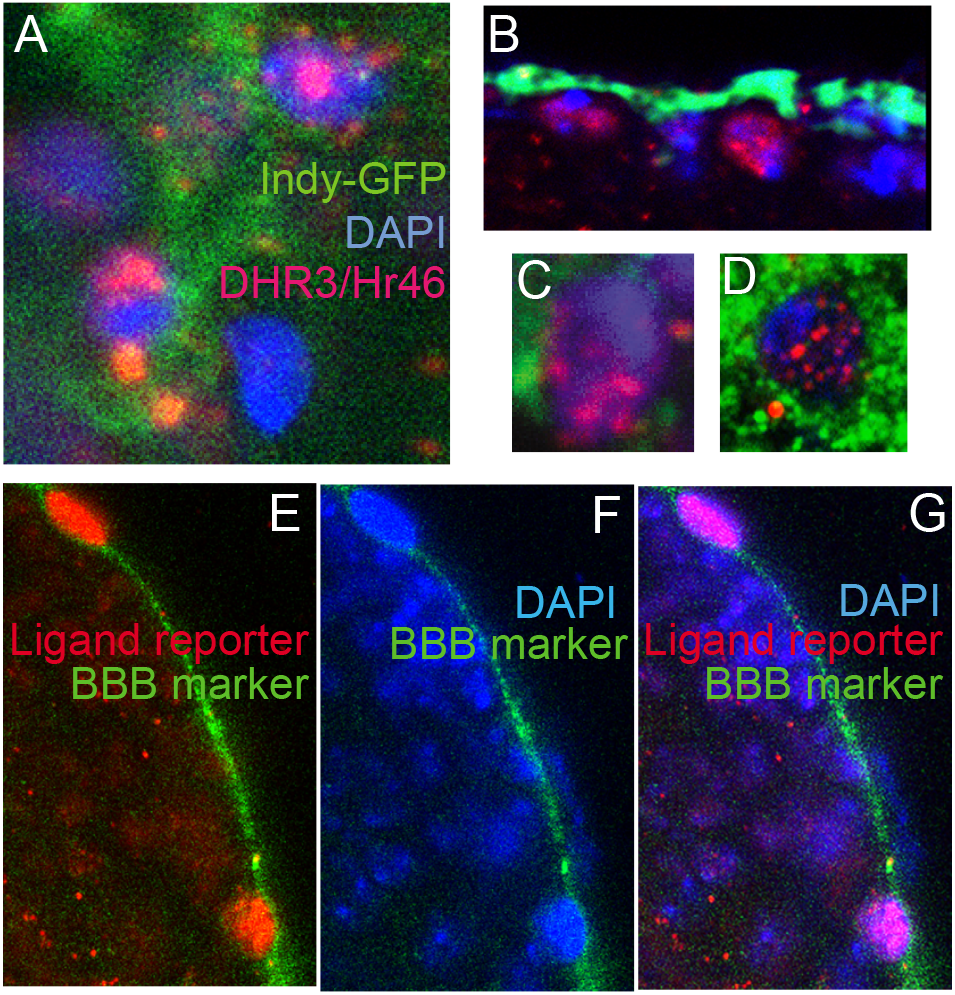
DHR3/Hr46 protein is located in SPG nuclei, and a reporter construct indicates that the DHR3/Hr46 ligand is present in the SPG cells of mature males. (A-D) Anti-DHR3 antibody staining (Red) shows the presence of DHR3 in the nuclei and cytoplasm of SPG cells. Indy-GFP (green) marks BBB cells. Blue: DNA staining (DAPI). (E-G) Activation of the *hs-Gal4^LBD^* ^DHR3^ reporter (25) indicates the presence of the DHR3/Hr46 ligand in SPG cells. *hs-Gal4^LBDDHR3^/UAS-dsRed/indy-GFP* mature males were heat-shocked to express *Gal4^LBDDHR3^.* Following binding of DHR3 ligand, Gal4 initiates transcription at *UAS-dsRed(nuclear).* dsRed can be seen expressed in the characteristic large nuclei of SPG cells (red). *Indy-GFP* expression is used as a BBB marker (green). Blue: DNA staining (DAPI, blue).

DHR3/Hr46 belongs to the family of ligand activated nuclear receptors. And while its ligand is unknown, insertion of the putative ligand binding domain into the Gal4 activation domain results in the transcriptional activation of Gal4 in cells containing the putative ligand. Palanker et al. have shown that this construct recapitulates Hr46 activation in cells where Hr46 transcriptional activity has been observed (25). Binding of the putative ligand activates *Gal4^LBD(DHR3)^* whose activity can then be visualized by a *UAS-reporter.* Importantly, in the construct, the *Gal4^LBD(DHR3)^* reporter is driven by a *hsp70* heat shock promoter *(hsp70-Gal4^LBD(DHR3^*). This makes it possible to interrogate the presence of the ligand at a time of choice. We induced the *Gal4^LBD(DHR3)^* ligand sensor in mature males by exposing the flies to 37 degrees Celsius for one hour and fixed their brains four hours later. We combined *hsp70-Gal4^LBD(DHR3^* with *UAS-dsRed* to visualize Gal4 activity, and *indy-GFP* for visualization of the BBB. As shown in Figs 4 E-G, dsRed staining is observed in the large nuclei of SPG cells, indicating that the ligand for Hr46 is present in these cells in adult mature males. Together our findings support a scenario in which Hr46 is activated and physiologically needed in the cells of adult males to support normal male courtship behavior.

## Discussion

Our microarray screen revealed that the *Drosophila* BBB contains male-enriched transcripts in males, as well as female-enriched transcripts in females. We have previously observed a reduction in male courtship when we conditionally feminized the BBB cells of mature males. Together these findings suggest that sex-specifically enriched transcripts contribute to a “male-specific” state of BBB cells that shapes its physiology and its dynamic interaction with the brain to modulate courtship. The previous feminization experiments were done in mature adult males by conditional induction of the female-specific TRA protein (26). TRA is a master controller of sex determination by virtue of its direct control of the two major sex specific transcription factors DSX and FRU, which in turn control a multitude of genes (27, 28). Non-induced males were normal, demonstrating that it was the acute adult change in transcripts that resulted in disturbed courtship. In the microarray experiments described here we sampled all transcripts that were present in the BBB cells of mature males and females. These animals were of the same age as the flies in the TRA induction studies. Neither males nor females had mating experience. The sex-specific transcripts we identify here therefore likely include transcripts that were affected in the feminization experiment.

We identified a total of 284 sex-preferentially expressed transcripts. It is likely that a number of them are required in the regulation of sex-specific behaviors and that their disruption will affect courtship. Identifying them holds the promise of insight into the physiological processes that underlie BBB-brain communication that is required for normal courtship. However, there will likely also be commonly expressed transcripts that participate in these sex-specific processes as they interact with sex-specific partners or regulators, or respond to sex-specific incoming signals. The majority of identified SPG transcripts are equally expressed in males and females, representing an insight into the overall transcriptional “makeup” of SPG cells of mature males and females. We expect many of them to overlap with the transcripts identified by deSalvo et al. (5). In contrast to our study, their characterization included both layers of the BBB, PG and SPG cells, without distinguishing between males and females.

DHR3/Hr46 belongs to the nuclear-receptor superfamily that is characterized by the presence of a highly conserved DNA-binding domain (DBD) and a less conserved C-terminal ligand-binding and dimerization domain (LBD). The ligand for DHR3/Hr46 is unknown, but the reporter construct made by Palanker et al. strongly indicates that a ligand exists that binds to the LBD in the receptor (25). In larvae, Palanker et al. observed fairly widespread, but not ubiquitous, activation, including in the fat body, leading to the speculation that DHR3 might have metabolic functions. ROR, the mammalian homologue of DHR3, is known to bind cholesterol and play a role in lipid homeostasis. Flies do not produce cholesterol, but take it up from their diet and it is an important precursor for the steroid ecdysone among other roles. Another nuclear receptor, DHR96 has been shown to bind cholesterol in *Drosophila* and to be essential for cholesterol homeostasis (29), but this does not exclude a role for DHR3/Hr46. Palanker et al. observed strong Gal4^DHR3LDB^ reporter expression in tissues of late third instars, with expression dropping in pre-pupariation, but strong activation was observed again in late pupae. We show here activation of the reporter construct in the BBB of mature adult males. In these experiments, the reporter construct is conditionally induced by a heat pulse in mature males. Thus, the observed activation reflects a “snapshot” of the presence of the putative ligand at that time. The observed activity coincides with the time when knockdown of DHR3 causes a reduction in courtship.

DH3/Hr46 is best known for its essential role in development as an ecdysone effector. It is activated by ecdysone and is a part of an activation cascade in response to ecdysone. It induces another nuclear receptor, Ftz-F1, among numerous other genes. Eventually, it acts as a negative feedback regulator to turn off ecdysone-receptor signaling (10–12, 30, 31). DHR3/Hr46 has essential functions during embryogenesis, prior to molts, and at the onset of metamorphosis. To our knowledge, this is the first report of an adult non-developmental role for DHR3/Hr46. Our conditional knockdown experiments demonstrate that its presence in the BBB of mature males is needed for normal courtship. Whether this reflects a role for an ecdysone-induced signaling cascade and transcriptional activation of downstream targets remains to be determined. Data from (32) suggest that ecdysone and the ecdysone receptor (EcR) are present in the BBB. We have likewise found in our screen that *EcR* and *ftz-F1* RNAs are present in SPG cells in a non-sex-specific manner. In analogy to its developmental role, DHR3/Hr46 most likely acts as a transcriptional regulator. Our observation that the Hr46 protein is present in SPG nuclei supports this interpretation. However, in an unexpected finding Montagne et al have identified DHR3 as a S6K interacting protein in late larvae/prepupae (33). Intriguingly, this function required a novel form of DHR3 that did not contain the DNA binding domain, but did require the ligand binding domain. The presence of this altered form of DHR3 increased phosphorylation activity of S6K. This finding, together with the short time scale of the response led the authors to propose an alternative non-genomic role of DHR3, possibly as a mediator of the metabolic state of these cells. We do not know whether this isoform plays a role in courtship and whether DHR3 might have a role that is independent of EcR in the BBB, conceivably in addition to the transcriptional role that is suggested by its presence in the nucleus.

Taken together, the data presented here demonstrate an adult physiological role for DHR3/Hr46, a nuclear receptor mainly known for its crucial function in development, in the glial cells of the BBB where it is required for the regulation of normal male courtship.

## Materials and Methods

### Fly stocks

*SPG-Gal4/TM3* (6) was a gift from Roland Bainton, UCSF. *tubP-Gal80^ts^/CyO* and *tubP-Gal80t^s^/TM3,Sb* flies were a gift from Gregg Roman (University of Mississippi). *DHR3/Hr46* RNAi lines *y^1^ v^1^;P{TRiP.JF02542}attP2* (BL 27253) and *y^1^ v^1^; P{TRiP.JFO2543}attP2* (BL 27254); *w^1118^; P{w[+mC]=UAS-RedStinger (dsRed)}4/CyO* (BL 8546); *w*; P{PTT-GC}IndyYC00l7/TM6C, Sb1 (Indy-GFP)* (BL 50860) were obtained from the Bloomington *Drosophila* stock center (https://bdsc.indiana.edu/). *y, repo-Gal4* on X was a gift from Takeshi Awasaki (University of Massachusetts (23); The*y* mutation was removed by recombination. *w; Pin, repo-Gal80/CyO* flies were a gift from Rob Jackson (Tufts University). Pin was removed by recombination. *w; +; repo-Gal80* flies (23) were a gift from Christian Klämbt (University of Münster). *w^1118^; P{w[+mC]=hs-GAL4-DHR3.LBD* was a gift from Carl Thummel (University of Utah).

### Gal80^ts^ experiments

*tubP-Gal80^ts^* carrying flies and control flies were raised at 18°C. Virgin males were collected at eclosion and kept in individual vials for 5-8 days at 18°C. Flies were then placed at 32°C for 24 hours for induction. Following induction, induced and uninduced flies were kept at 25°C overnight prior to courtship assays.

### Behavioral assays

The courtship assay and activity assay were performed as previously described (34). In short, males were placed in a plexiglass “mating wheel” (diameter 0.8 cm), together with a 2-4 hrs old *Canton-S* virgin female. The courtship index was calculated as the fraction of time the male spent displaying any element of courtship behavior (orienting, following, wing extension, licking, attempted copulation, copulation) within a 10-minute observation period (18). Short-term activity assays were performed as previously described (22). Individual males were placed into the “mating wheel” containing a filter paper with a single line dividing the chamber in half. After 2-3 minutes of acclimation time, the number of times the male crossed the center line within the three-minute observation time was counted.

Each graph represents sets of control and experimental genotypes that were grown, collected, aged and tested in parallel. In each behavioral session, equal numbers of all genotypes were tested.

### Microarray analysis

To isolate blood-brain barrier cells, flies bearing the *SPG-Gal4* driver were crossed to flies carrying the fluorescent reporter transgene, *UAS-DsRed*. This resulted in the expression of the visible fluorescent marker DsRed to mark the nuclei of BBB cells. Prior to the experiment, both the driver *SPG-Gal4* and the *UAS-DsRed* lines were outcrossed with a Cantonized *w^1118^* strain for 10 generation. The flies were grown in a 25°C incubator under a 12hrs light/12hrs dark cycle. Eclosing males and females were collected and kept in separate groups of 10-15 flies of the same sex under the same conditions for 4 days and then dissected between ZT 5 and ZT 7 to control for levels of cycling transcripts. Equal numbers of males and females originating from the same culture were dissected in each sitting. The brains were dissected in icecold 1 X PBS. The dissected brains were then transferred to a new petri dish containing ice-cold 1X PBS within half an hour. Carefully, under the fluorescent microscope, individual and/or groups of blood-brain barrier cells marked with DsRed were isolated manually by using Dumont # 5 SF superfine forceps (Fine Science Tools, Inc). The cells were then immediately transferred to a frozen droplet of Trizol reagent on dry ice and stored in −80°C until further processing. Cells were isolated from at least 50 brains for each genotype. The approximate total number of cells isolated per brain varied from ~60-120. The forceps were cleaned with RNAZap when moving from one genotype to the other.

The isolated BBB cells of male and female flies were provided to GenUs Biosystems (http://www.genusbiosystems.com/) for microarray analysis. A total of 3 biological replicates for males and females were submitted. Cells were lysed in TRI reagent (Ambion) and Total RNA was isolated using phenol/chloroform extraction followed by purification over RNeasy spin columns (Qiagen). The concentration and purity of Total RNA was measured by spectrophotometry at OD260/280 and the quality of the Total RNA sample was assessed using an Agilent Bioanalyzer with the RNA6000 Nano Lab Chip (Agilent Technologies). Labeled cRNA was prepared by linear amplification of the Poly (A)+ RNA population within the Total RNA sample. Briefly, 1 μg of Total RNA was reverse transcribed after priming with a DNA oligonucleotide containing the T7 RNA polymerase promoter 5’ to a dT(24) sequence. After second-strand cDNA synthesis and purification of double-stranded cDNA, in vitro transcription was performed using T7 RNA polymerase. The quantity and quality of the cRNA was assayed by spectrophotometry and on the Agilent Bioanalyzer. One microgram of purified cRNA was fragmented to uniform size and applied to *Drosophila* (V2) Gene Expression microarray (Agilent Technologies, Design ID 021791) in hybridization buffer. Arrays were hybridized at 37° C for 18 hrs in a rotating incubator, washed, and scanned with a G2565 Microarray Scanner (Agilent Technologies). Arrays were processed with Agilent Feature Extraction software and data was analyzed with GeneSpring GX software (both Agilent Technologies). To compare individual expression values across arrays, raw intensity data from each gene was normalized to the 75th percentile intensity of each array. Genes with values greater than background intensity in all female or all male replicates were filtered for further analysis. Differentially expressed genes were identified with fold change > 2-fold and Welch Ttest, p-value < 0.05.

The data discussed in this publication have been deposited in NCBI’s Gene Expression Omnibus and are accessible through GEO Series accession number GSE 157122 (https://www.ncbi.nlm.nih.gov/geo/query/acc.cgi?acc=GSE157122).

### Generation of Mdr65-Gal4 transgenic flies

650 bp of sequence upstream of the *Mdr65* coding sequence was PCR amplified from *CS* genomic DNA using the primers *5’cggaattc(EcoRI)TCCATCACTTAGCAAAGCAGACTTCAATC* and 5’cgggatcc(BamH1) *GGTGATGTTTAGTCGGCACTGACGA* and inserted into the *Drosophila* transformation vector *pGATN* to create *Mdr65-Gal4.* In *pGATN,* expression of the yeast transcription factor Gal4 is driven by the inserted promoter sequences (13). Transgenic flies were generated by *Rainbow Transgenic Flies* (https://www.rainbowgene.com/) by P-element mediated insertion. The expression pattern in *Mdr65-Gal4* transgenic lines was examined by crosses to *UAS-dsRed* (nuclear) or *UAS-mcD8-dsRed.*

### Immunohistochemistry

Immunohistochemistry on isolated brains was performed as described in Li et al. (35). The DHR3/Hr46 antibody was a gift from Carl Thummel, University of Utah (11) and was used at 1:50 dilution. To visualize BBB cells, flies carrying *Indy-GFP* were used for anti-DHR3/ anti-GFP double staining. *Indy-GFP* marks BBB cells (36). Antibodies used:

Rabbit anti-DHR3, 1:50 (gift of Carl Thummel, University of Utah (11)); Rabbit anti-RFP (abcam ab62341), 1:200; chicken anti-GFP (abcam ab13970), 1:500; Alexa Fluor 555 goat anti-rabbit (Invitrogen A21429) 1:200; Alexa Fluor 635 goat antimouse (Invitrogen A31575) 1:200; Alexa Fluor 488 goat anti-chicken (Thermo Fisher Scientific A-11039).

Injection of 10kd Dextran-TR to assess the integrity of the BBB was performed as described in Hoxha et al. (8).

### Test for presence of DHR3/Hr46 ligand

*hs-Gal4^LBD(DHR3)^* flies were crossed to *UAS-dsRed(nuclear); indy-GFP* flies. Progeny were collected at eclosion and kept in small groups of males or females for 4 days. Expression of *hs-Gal4^LBD(DHR3^* was induced by placing flies in prewarmed food vials at 37°C for one hour, followed by recovery at room temperature for three hours and brain isolation. dsRed expression as a measure of Gal4 activation by DHR3 ligand and GFP (as BBB marker) were assessed by immunohistochemistry.

### Statistical Analysis

Two-way analysis of variance (ANOVA) was used to establish overall significance. Post hoc analysis for multiple comparisons was carried out with Tukey (HSD). P values < 0.05 were considered statistically significant. All statistical calculations were done using XLSTAT (Addinsoft, NY, NY) running on Microsoft Excel for Mac (version 16). All ±error bars are standard error of the mean (SEM).

---

**S1 Table.**
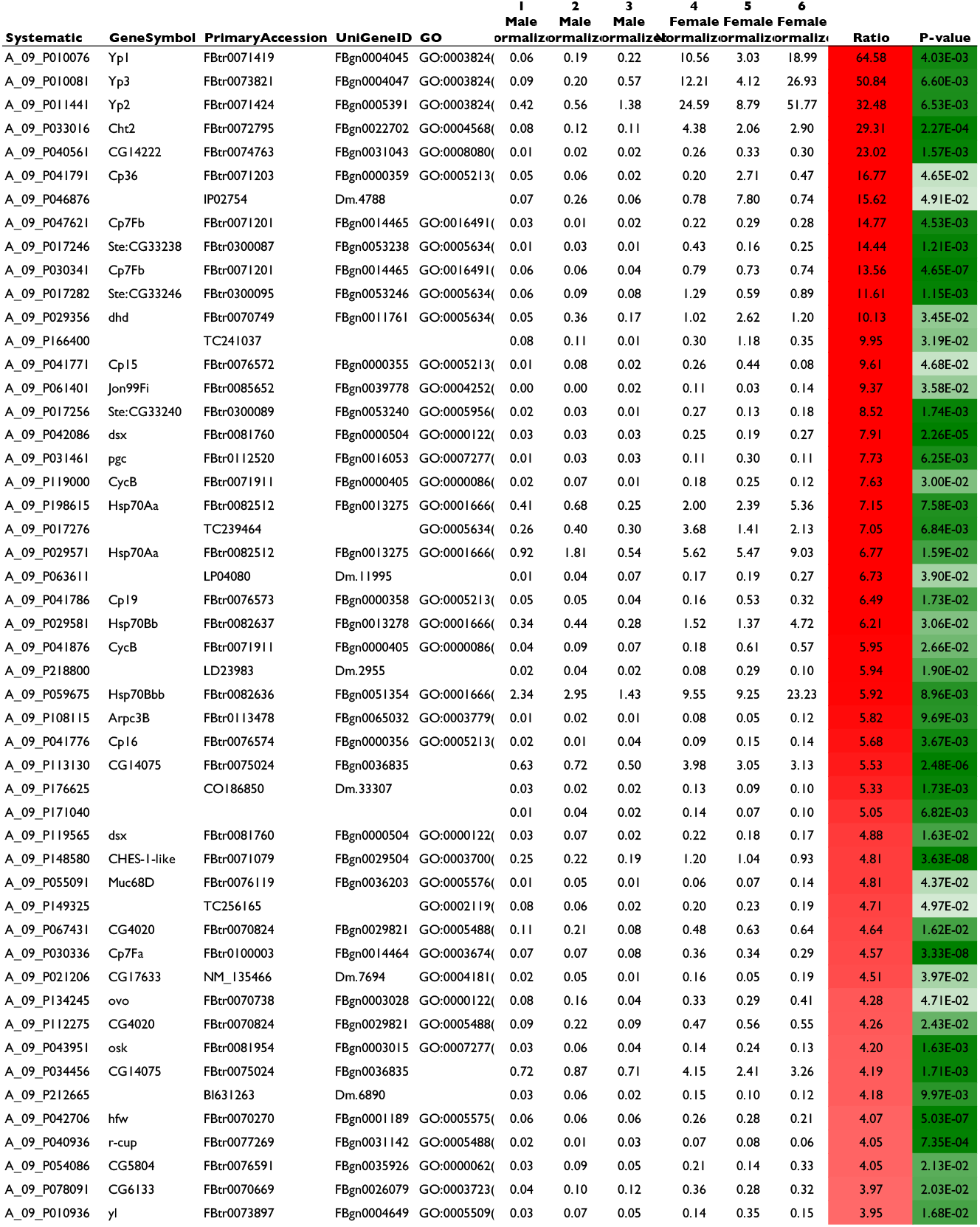
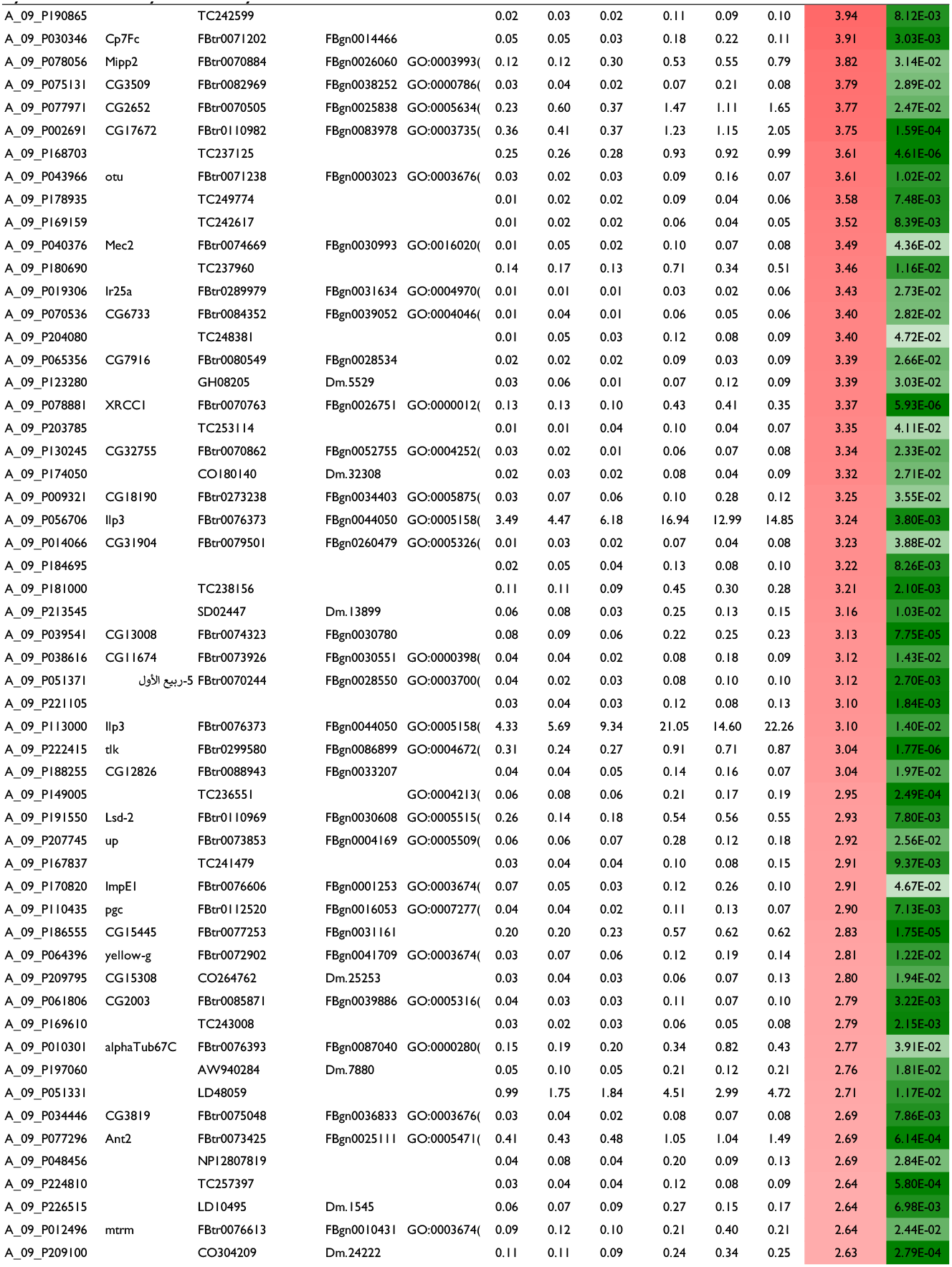
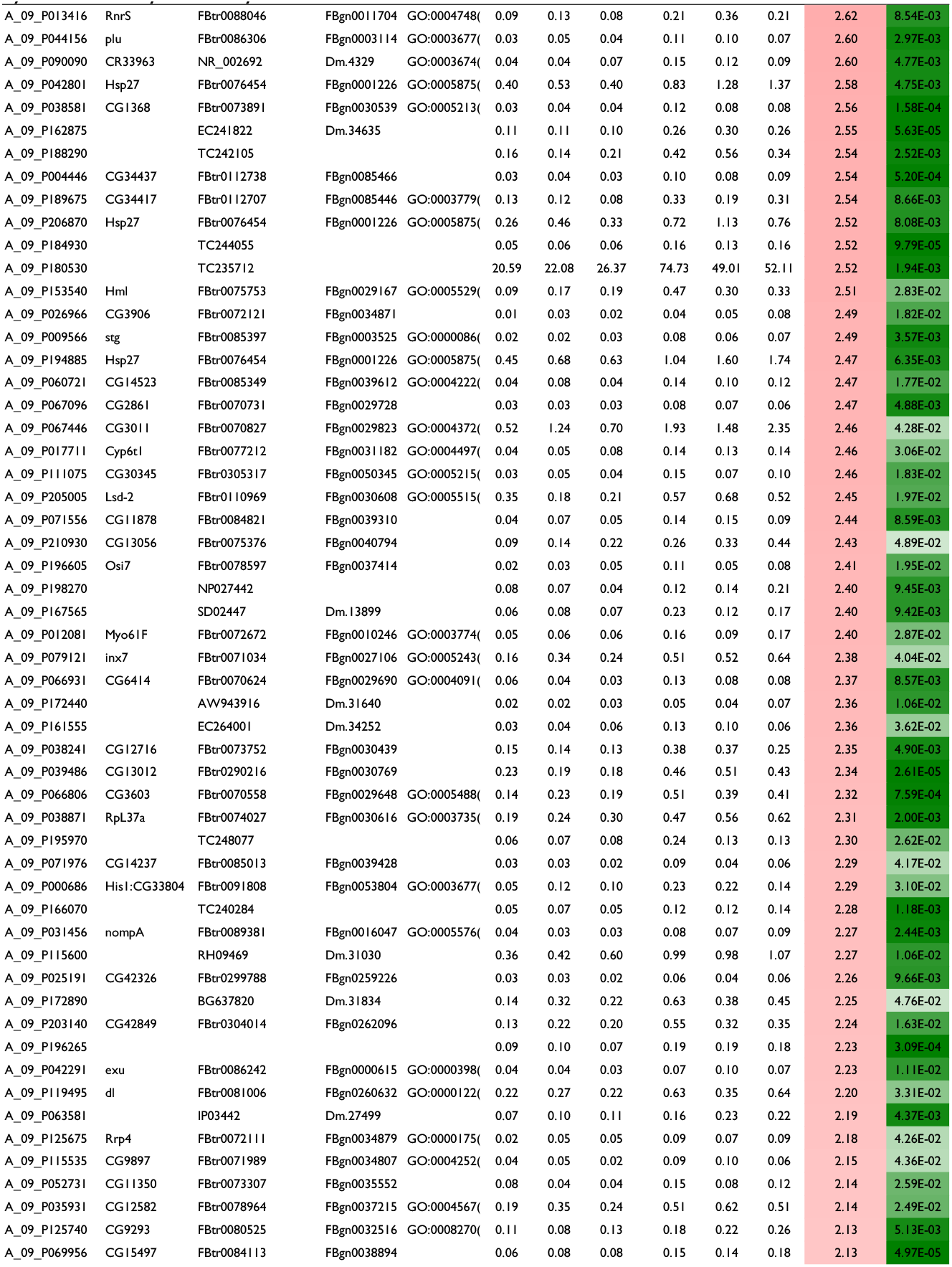
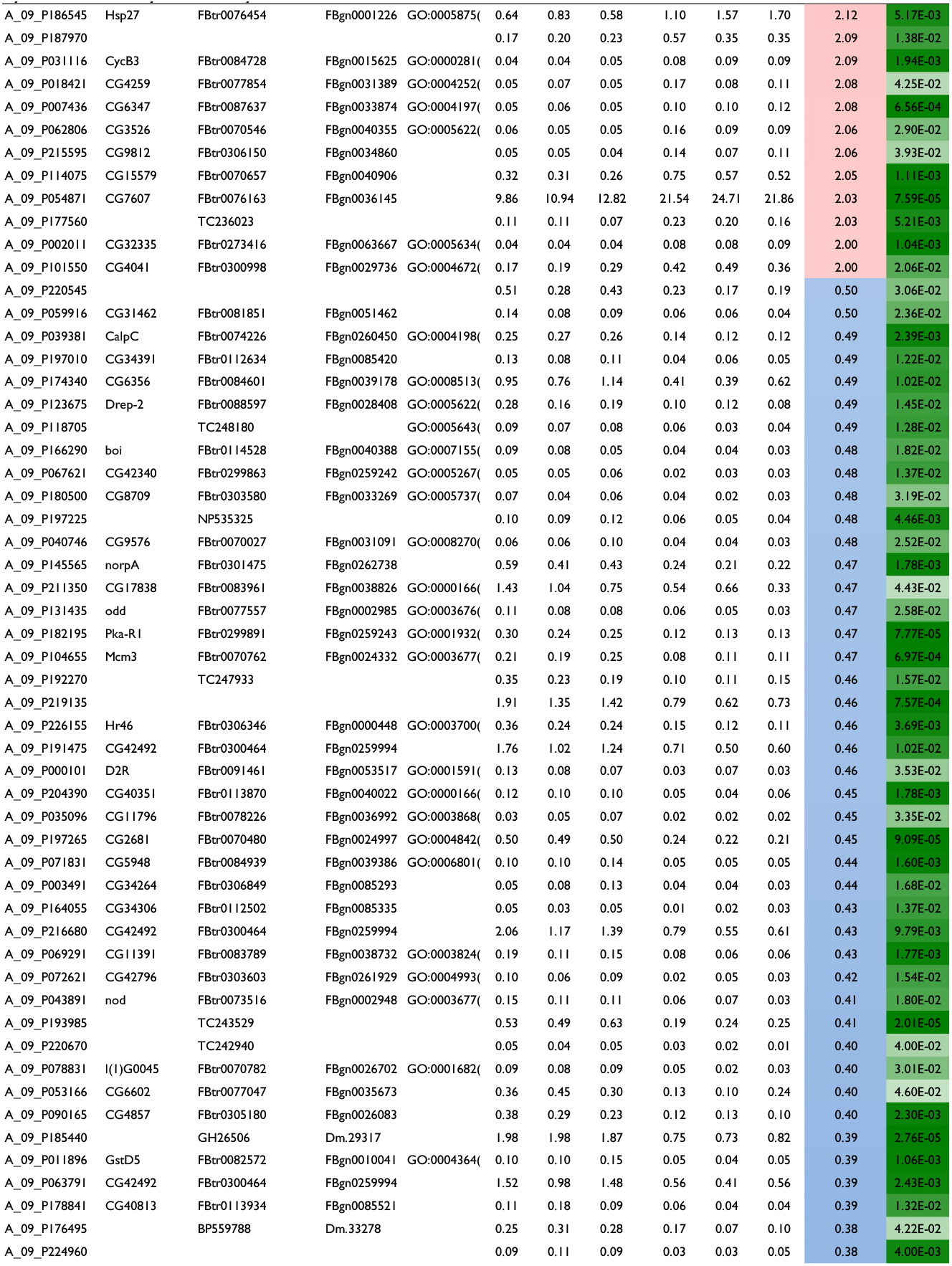
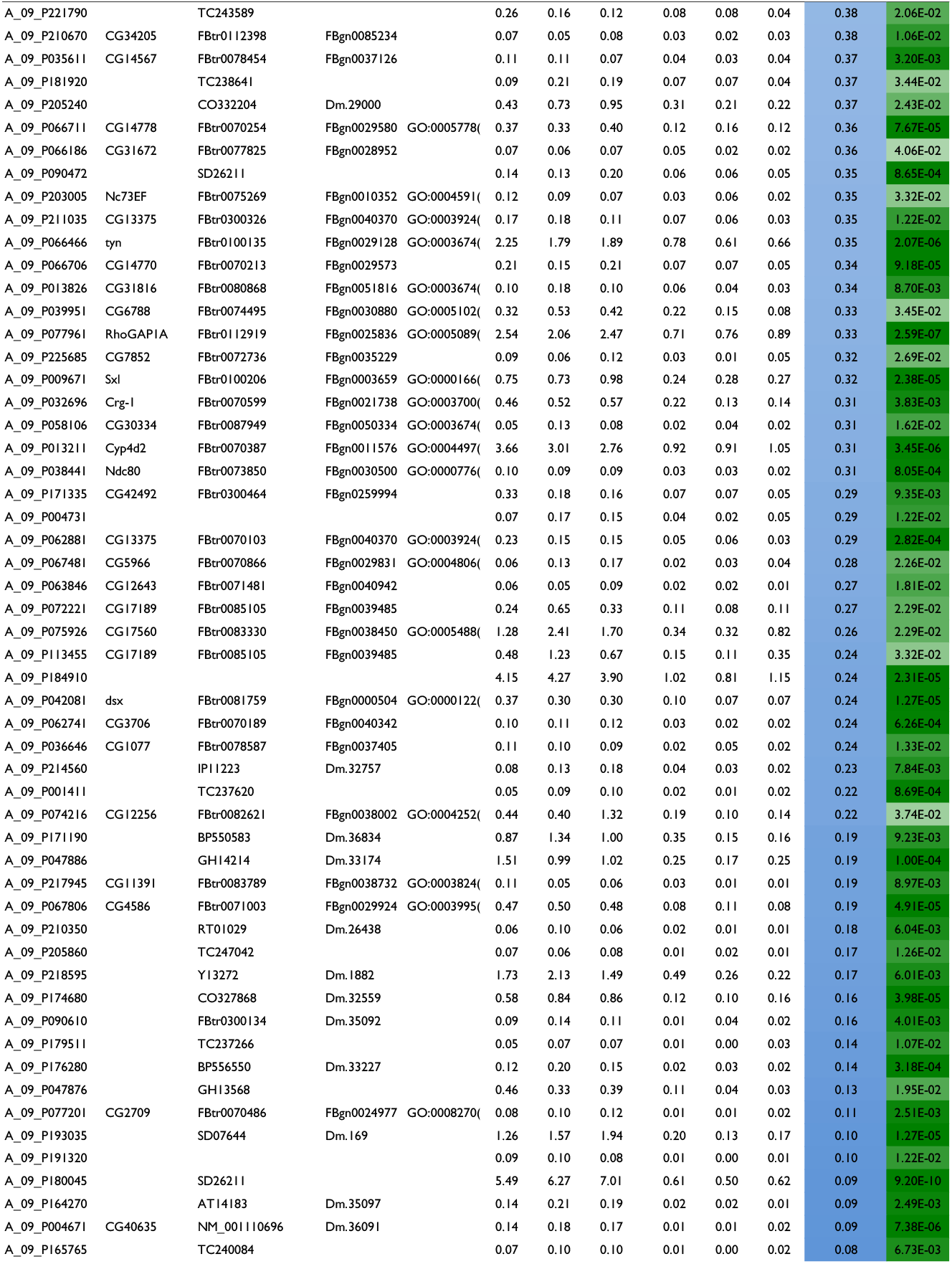
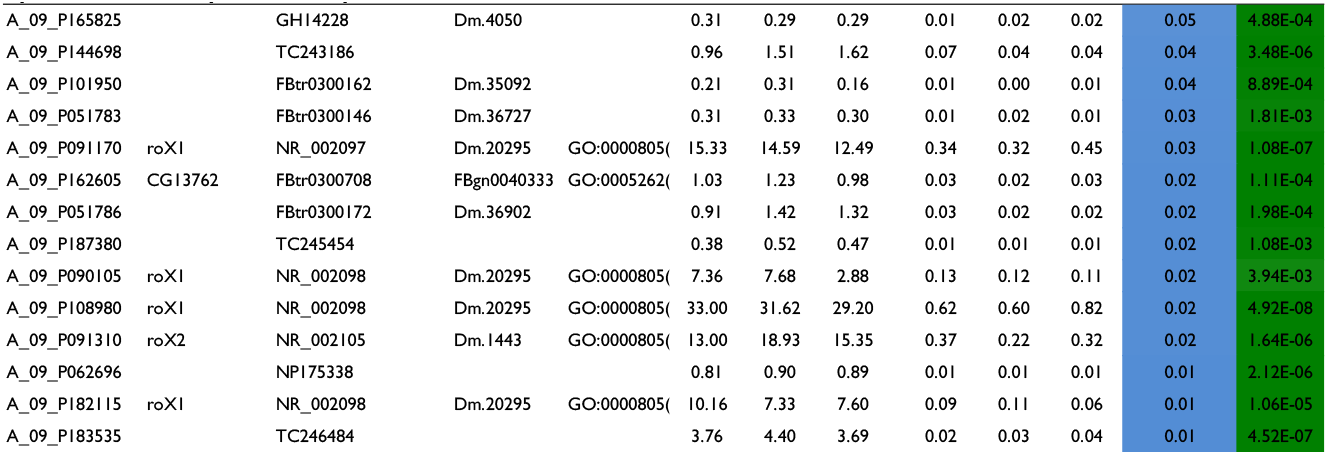
Differentially expressed genes in BBB from males vs. females (>2Fold, p<0.05). 284 transcripts were identified that differ by >2Fold (p<0.05). Of those 112 were male-preferentially expressed, 172 were female-preferentially expressed. Intensity values are normalized to the 75th percentile intensity of each array. p-values are based on a Welch T-test.

**S2 Table.** Gene Ontology classifications of the 284 genes differentially expressed in BBB Males vs. Females. Category=the name of the category within the ontology. Genes in Category=the total number of genes in the genome that have been assigned to the category Genes in List in Category=the total number of genes that are both in the selected gene list and in the category. P-value=a hypergeometric p-value without multiple testing corrections.

| Biological Process |  |  |  |
| --- | --- | --- | --- |
| Category | Genes in Category | Genes in List in Category | p-Value |
| GO:16457: dosage compensation complex assembly (sensu Insecta) | 13 | 5 | 6.77E-08 |
| GO:42714: dosage compensation complex assembly | 13 | 5 | 6.77E-08 |
| GO:30237: female sex determination | 28 | 6 | 1.55E-07 |
| GO:35079: polytene chromosome puffing | 7 | 4 | 2.18E-07 |
| GO:35080: heat shock-mediated polytene chromosome puffing | 7 | 4 | 2.18E-07 |
| GO:9047: dosage compensation, by hyperactivation of X chromosome | 22 | 5 | 1.30E-06 |
| GO:7549: dosage compensation | 48 | 6 | 4.36E-06 |
| GO:42026: protein refolding | 15 | 4 | 8.05E-06 |
| GO:7296: vitellogenesis | 16 | 4 | 1.07E-05 |
| GO:19102: male somatic sex determination | 6 | 3 | 1.41E-05 |
| GO:45496: male analia morphogenesis (sensu Endopterygota) | 6 | 3 | 1.41E-05 |
| GO:45497: female analia morphogenesis (sensu Endopterygota) | 6 | 3 | 1.41E-05 |
| GO:7530: sex determination | 62 | 6 | 1.97E-05 |
| GO:9408: response to heat | 128 | 8 | 2.06E-05 |
| GO:51084: posttranslational protein folding | 19 | 4 | 2.22E-05 |
| GO:48086: negative regulation of pigmentation | 7 | 3 | 2.46E-05 |
| GO:19101: female somatic sex determination | 21 | 4 | 3.38E-05 |
| GO:30238: male sex determination | 8 | 3 | 3.91E-05 |
| GO:281: cytokinesis after mitosis | 8 | 3 | 3.91E-05 |
| GO:9266: response to temperature stimulus | 144 | 8 | 4.81E-05 |
| GO:48071: sex-specific pigmentation | 12 | 3 | 1.50E-04 |
| GO:7486: female genitalia morphogenesis (sensu Endopterygota) | 13 | 3 | 1.93E-04 |
| GO:30540: female genitalia morphogenesis | 13 | 3 | 1.93E-04 |
| GO:7487: analia morphogenesis (sensu Endopterygota) | 13 | 3 | 1.93E-04 |
| GO:7079: mitotic chromosome movement towards spindle pole | 3 | 2 | 2.42E-04 |
| GO:7485: male genitalia morphogenesis (sensu Endopterygota) | 15 | 3 | 3.03E-04 |
| GO:35263: genital disc sexually dimorphic development | 15 | 3 | 3.03E-04 |
| GO:45498: sex comb development | 15 | 3 | 3.03E-04 |
| GO:45934: negative regulation of nucleobase, nucleoside, nucleotide and nucleic acid metabolism | 348 | 11 | 3.15E-04 |
| GO:18993: somatic sex determination | 39 | 4 | 4.10E-04 |
| GO:19099: female germ-line sex determination | 17 | 3 | 4.47E-04 |
| GO:18992: germ-line sex determination | 18 | 3 | 5.33E-04 |
| GO:1666: response to hypoxia | 45 | 4 | 7.12E-04 |
| GO:7484: genitalia morphogenesis (sensu Endopterygota) | 20 | 3 | 7.35E-04 |
| GO:19953: sexual reproduction | 1413 | 25 | 7.67E-04 |
| GO:92: mitotic anaphase B | 5 | 2 | 7.96E-04 |
| GO:7548: sex differentiation | 168 | 7 | 8.28E-04 |
| GO:8608: attachment of spindle microtubules to kinetochore | 6 | 2 | 1.19E-03 |
| GO:51305: chromosome movement towards spindle pole | 6 | 2 | 1.19E-03 |
| GO:31324: negative regulation of cellular metabolism | 420 | 11 | 1.48E-03 |
| GO:50832: defense response to fungi | 56 | 4 | 1.63E-03 |
| GO:45944: positive regulation of transcription from RNA polymerase II promoter | 96 | 5 | 1.77E-03 |
| GO:86: G2/M transition of mitotic cell cycle | 27 | 3 | 1.80E-03 |
| GO:9620: response to fungi | 59 | 4 | 1.97E-03 |
| GO:16481: negative regulation of transcription | 312 | 9 | 2.10E-03 |
| GO:45892: negative regulation of transcription, DNA-dependent | 312 | 9 | 2.10E-03 |
| GO:7483: genital disc morphogenesis | 29 | 3 | 2.22E-03 |
| GO:48070: regulation of developmental pigmentation | 30 | 3 | 2.45E-03 |
| GO:30730: sequestering of triacylglycerol | 9 | 2 | 2.80E-03 |
| GO:9892: negative regulation of metabolism | 458 | 11 | 2.92E-03 |
| GO:7276: gametogenesis | 1385 | 23 | 2.93E-03 |
| GO:7277: pole cell development | 33 | 3 | 3.23E-03 |
| GO:50000: chromosome localization | 33 | 3 | 3.23E-03 |
| GO:51303: establishment of chromosome localization | 33 | 3 | 3.23E-03 |
| GO:3: reproduction | 1571 | 25 | 3.35E-03 |
| GO:45570: regulation of imaginal disc growth | 36 | 3 | 4.14E-03 |
| GO:31887: lipid particle transport along microtubule | 11 | 2 | 4.23E-03 |
| GO:30539: male genitalia morphogenesis | 37 | 3 | 4.48E-03 |
| GO:51338: regulation of transferase activity | 74 | 4 | 4.50E-03 |
| GO:45859: regulation of protein kinase activity | 74 | 4 | 4.50E-03 |
| GO:45448: mitotic cell cycle, embryonic | 80 | 4 | 5.93E-03 |
| GO:7446: imaginal disc growth | 82 | 4 | 6.47E-03 |
| GO:6950: response to stress | 1055 | 18 | 6.59E-03 |
| GO:51656: establishment of organelle localization | 132 | 5 | 6.92E-03 |
| GO:122: negative regulation of transcription from RNA polymerase II promoter | 133 | 5 | 7.14E-03 |
| GO:16545: male courtship behavior (sensu Insecta), wing vibration | 44 | 3 | 7.29E-03 |
| GO:45433: male courtship behavior (sensu Insecta), song production | 44 | 3 | 7.29E-03 |
| GO:46660: female sex differentiation | 44 | 3 | 7.29E-03 |
| GO:16542: male courtship behavior (sensu Insecta) | 45 | 3 | 7.76E-03 |
| GO:48065: male courtship behavior (sensu Insecta), wing extension | 45 | 3 | 7.76E-03 |
| GO:35186: syncytial blastoderm mitotic cell cycle | 45 | 3 | 7.76E-03 |
| GO:42742: defense response to bacteria | 142 | 5 | 9.34E-03 |
| **Cellular Component** |  |  |  |
| **Category** | **Genes in Category** | **Genes in List in Category** | **p-Value** |
| GO:805: X chromosome | 18 | 5 | 5.19E-07 |
| GO:803: sex chromosome | 20 | 5 | 9.26E-07 |
| GO:46536: dosage compensation complex | 21 | 5 | 1.21E-06 |
| GO:16456: dosage compensation complex (sensu Insecta) | 21 | 5 | 1.21E-06 |
| GO:5956: protein kinase CK2 complex | 15 | 4 | 9.29E-06 |
| GO:45495: pole plasm | 31 | 5 | 9.34E-06 |
| GO:43232: intracellular non-membrane-bound organelle | 2013 | 37 | 2.31E-05 |
| GO:43228: non-membrane-bound organelle | 2013 | 37 | 2.31E-05 |
| GO:15630: microtubule cytoskeleton | 874 | 20 | 1.68E-04 |
| GO:5856: cytoskeleton | 1120 | 23 | 2.56E-04 |
| GO:5875: microtubule associated complex | 700 | 17 | 2.76E-04 |
| GO:43229: intracellular organelle | 5737 | 72 | 5.01E-04 |
| GO:43226: organelle | 5737 | 72 | 5.01E-04 |
| GO:42600: chorion | 20 | 3 | 8.17E-04 |
| GO:43234: protein complex | 3209 | 46 | 8.38E-04 |
| GO:43186: P granule | 22 | 3 | 1.09E-03 |
| GO:5622: intracellular | 7566 | 86 | 2.42E-03 |
| GO:19908: nuclear cyclin-dependent protein kinase holoenzyme complex | 13 | 2 | 6.37E-03 |
| GO:307: cyclin-dependent protein kinase holoenzyme complex | 14 | 2 | 7.38E-03 |
| GO:30312: external encapsulating structure | 45 | 3 | 8.58E-03 |
| **Molecular Function** |  |  |  |
| **Category** | **Genes in Category** | **Genes in List in Category** | **p-Value** |
| GO:5213: structural constituent of chorion (sensu Insecta) | 10 | 5 | 9.76E-09 |
| GO:19887: protein kinase regulator activity | 93 | 8 | 1.18E-06 |
| GO:19207: kinase regulator activity | 97 | 8 | 1.62E-06 |
| GO:16903: oxidoreductase activity, acting on the aldehyde or oxo group of donors | 63 | 4 | 1.98E-03 |
| GO:16538: cyclin-dependent protein kinase regulator activity | 38 | 3 | 4.02E-03 |
| GO:5198: structural molecule activity | 843 | 15 | 5.06E-03 |
| GO:15245: fatty acid transporter activity | 13 | 2 | 5.21E-03 |
| GO:5324: long-chain fatty acid transporter activity | 13 | 2 | 5.21E-03 |
| GO:16620: oxidoreductase activity, acting on the aldehyde or oxo group of donors, NAD or NADP as acceptor | 48 | 3 | 7.75E-03 |

## Acknowledgements

We thank Dr. Carl Thummel for anti-DHR3 antibodies and fly lines and Takeshi Awasaki (University of Massachusetts), Rob Jackson (Tufts University), Christian Klämbt (University of Münster) and Gregg Roman (University of Mississippi) for fly lines. We thank Calvin Do for help with brain dissections. We thank GenUS biosystems for expert help with the microarray experiments.

